# MITF is essential for autophagy in the retinal pigment epithelium

**DOI:** 10.64898/2026.05.22.727222

**Authors:** Andrea García-Llorca, Kristján Hermannsson, Filippo Locri, Helder André, Eiríkur Steingrimsson, Margrét Helga Ogmundsdottir, Thor Eysteinsson

## Abstract

The Microphthalmia-associated transcription factor (MITF) plays a critical role in retinal pigment epithelium (RPE) development and function. Dysfunctional autophagy and lysosomal degradation in the RPE have been implicated in age-related retinal degeneration, yet the contribution of MITF to these pathways remains incompletely understood. Here, we show that reduced *Mitf* expression impairs autophagy in mouse and human RPE cells. Primary RPE cells from *Mitf*^*mi-vga9/+*^ heterozygotes mice displayed altered autophagic flux characterized by accumulation of LC3B-II and p62, while MITF knockdown in human ARPE-19 cells promoted autophagosome accumulation. Ultrastructural analysis further revealed age-dependent accumulation of autolysosomes and lipofuscin-like granules in mutant RPE cells. In addition, expression of autophagy-related genes was altered in mutant RPE tissue, supporting disrupted lysosomal-autophagic homeostasis. Together, our findings identify MITF as an important regulator of autophagy in the RPE and suggest that impaired MITF-dependent homeostasis may contribute to retinal degeneration.

## Introduction

The retinal pigment epithelium (RPE) of the eye serves several functions that are of importance to the viability of the retinal photoreceptors and secondary neurons. One of these functions is the degradation of debris from the photoreceptors via autophagy. This mechanism is indispensable for maintaining the internal stability of cells and protecting them from damage and disease^1–4^. In addition, autophagy is involved in many biological processes, including growth, differentiation, maintenance and remodeling of cells, and its disruption has been shown to be a factor in a range of human health disorders, including cancers, neurodegeneration, heart conditions and pathogen infections^5–8^. High baseline autophagy activity is exhibited by a healthy RPE. This is understandable, as these cells dwell in adverse surroundings, particularly for a cellular type intended to persist over a lifetime^9,10^. Apart from the considerable strain exerted on them by phagocytosis of photoreceptor outer segments, RPE cells are further tasked with handling continuous light exposure over their lifespan, elevated metabolic functions and encountering one of the highest oxygen levels found in the body^11^. Each of these aspects contributes to the significant vulnerability of the RPE to oxidative stress and the buildup of waste. The process of autophagy is essential in post-mitotic cells like the RPE for maintaining homeostasis and avoiding the progressive aggregation of impaired or deleterious constituents.

The microphthalmia-associated transcription factor family (MiT family) is composed of MITF, TFEB (transcription factor EB), TFE3 (transcription factor E3) and TFEC (transcription factor EC). It has been reported that TFEB and TFE3 are involved in the regulation of expression of genes involved in lysosome formation and autophagy induction in many cell types^12,13^. In addition, it has been recently shown that MITF plays an important role in regulating starvation induction in melanoma cells^14^. MITF, which is encoded by the *Mitf* gene, can bind to certain DNA sequences in the regulatory regions of a wide range of target genes ^15–17^. All mice with mutations in the *Mitf* gene share faulty neural-crest-derived melanocytes, resulting in lack of melanocytes in skin, eye, inner ear and other tissues. Furthermore, the mice exhibit loss of mast cells^18^ and problems with secondary bone resorption^19,20^. It has been demonstrated that the number of retinal vessels and their diameter are significantly different between *Mitf* mutant and control mice^21^. Human MITF mutations have been reported in individuals with the hearing loss and pigmentation disorders, Tietz and Waardenburg syndromes and they have been linked to microphthalmia and osteopetrosis^22,23^. Many processes that are essential to the integrity and functionality of photoreceptors are controlled and regulated by MITF in the RPE, such as the visual cycle, melanin formation, cell migration, antioxidant activities, proliferation, growth factor secretion, and RPE development and differentiation^3,24–26^. It has been proposed that MITF regulates lysosomal activity and autophagy through transcriptional control of autophagosomal and lysosomal genes. This regulatory function has been particularly well studied in melanoma cells ^17,27^. However, the role of *Mitf* in regulating autophagy in RPE has not been investigated before. The *Mitf*^*mi-vga9*^ mutation is a transgene insertion mutation that disrupts normal *Mitf* expression and results in ocular hypopigmentation and retinal abnormalities^28,29^. Homozygous *Mitf*^*mi-vga9*^ mice exhibit severe microphthalmia and profound retinal defects, whereas heterozygous animals display milder phenotypes while retaining overall retinal structure and viability^29^. For this reason, heterozygous *Mitf*^*mi-vga9/+*^ mice were used in this study to investigate the effects of reduced Mitf function on RPE autophagy and lysosomal homeostasis. Here, we show that the autophagy process is altered in RPE cells of mice heterozygous for the *Mitf*^*mi-vga9/+*^ mutation, indicating a role of *Mitf* in autophagy and lipofuscin-like granule formation/accumulation in RPE cells. The increase observed in the autophagy marker LC3B may reflect accumulation of autophagosomes due to impaired degradation rather than enhanced autophagic activity, suggesting that overall autophagy flux may still be reduced in the aging RPE.

## Results

### Gene expression analysis

To investigate the role of *Mitf* in autophagy in RPE cells, we determined the expression of the *Mitf, Lc3b, Lamp1* and *Rab7* genes by RT-PCR in eyecup tissues from wild type and *Mitf*^*mi-vga9/+*^ heterozygous mice at 3 and 12 months of age (N= 6 animals per genotype, both eyecups from each animal were pooled for analysis). *Mitf* expression was reduced in the mutant compared to control animals in all age groups studied (Figure 1). Due to the nature of the *Mitf*^*mi-vga9/+*^ mutation these mice are expected to express *Mitf* only from the wild type allele^28^, which would predict approximately 50% mRNA expression levels. However, we observed an approximately 80% reduction in *Mitf* expression, suggesting that expression from the remaining wild type allele may also be reduced. These findings indicate that *Mitf* expression in *Mitf*^*mi-vga9/+*^ mice may not reflect a simple gene dosage effect. The expression of *Lc3B* expression was increased in 3-month-old *Mitf*^*mi-vga9/+*^ heterozygotes compared to age-matched wild type but was reduced in 12-month-old *Mitf*^*mi-vga9/+*^ heterozygotes (Figure 1). Such an age-dependent decline may indicate that the early increase in *Lc3b* is not sustained over time, possibly reflecting a failure of compensatory mechanisms and progressive age-related dysregulation of the autophagy-lysosome system in aging *Mitf*^*mi-vga9/+*^ mice. The expression of *Lamp1* was reduced in the *Mitf*^*mi-vga9/+*^ heterozygotes regardless of age. The expression of the late endosomal gene *Rab7* was increased in *Mitf*^*mi-vga9/+*^ heterozygotes compared to wild type in all age groups studied. Together, these results suggest that both autophagy and the late endosomal pathways are altered in eyes of *Mitf*^*mi-vga9/+*^ heterozygotes.

**Figure 1.**
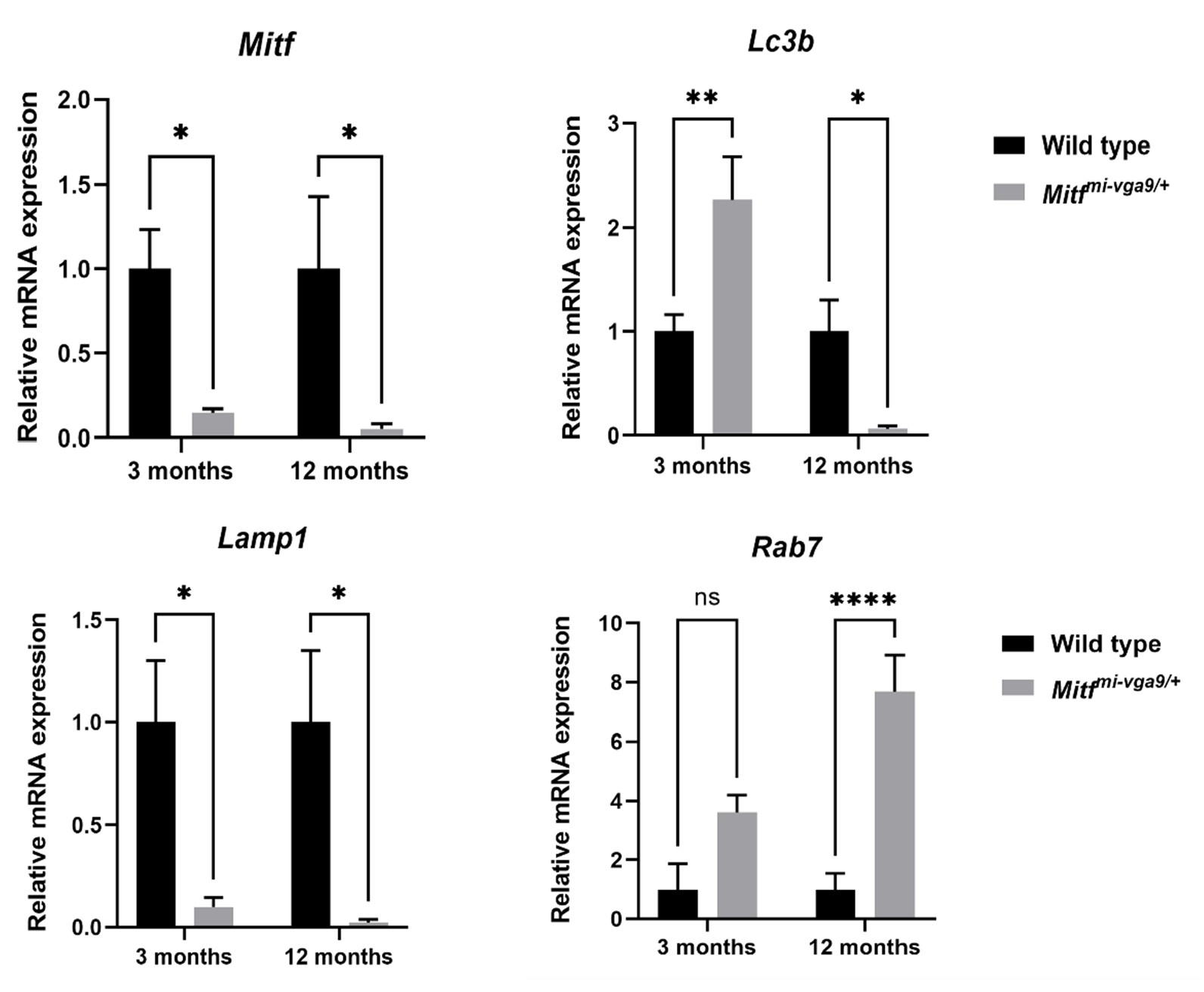
Quantitative RT-PCR analysis of eyecups from 3- and 12-month-old wild type and *Mitf*^*mi-vga9/+*^ mouse eyes. Primers specific for *Mitf, Lc3b, Lamp1* and *Rab7* were used to quantify the respective RNAs. *P<0.05, **P<0.01, ***P<0.001. The values on the graphs are mean ± SEM. P-values were calculated using the two-way ANOVA followed by Šidák post hoc multiple comparisons test. N=6 animals per genotype. Values are shown as relative mRNA expression normalized to *TBP* and *HPRT1* and expressed relative to age-matched controls.

### Lipofuscin-like granules accumulate during aging in *Mitf*^*mi-vga9/+*^ mice

We have previously analyzed fundus images from eyes of 3-month-old *Mitf*^*mi-vga9/+*^ and wild type mice^29^. Distinct yellow lesions were visible in the fundi of *Mitf*^*mi-vga9/+*^ mice, together with slight general hypopigmentation and a fine scattering of pigment mottling throughout the complete fundus. Representative bright field fundus photographs from the eyes of wild type and *Mitf*^*mi-vga9/+*^ heterozygotes at 6 and 12 months of age (n=6) were obtained with a fundus camera (Figure 2). The normal size of the optic disk areas in *Mitf*^*mi-vga9/+*^ mice, evident from the bright field images, confirmed that the eyes of this mutant grow to a normal size at 6- and 12-months-old compared to age-matched wild type mice (Figure 2). Fundus images from the mutant showed yellow lesions over the entire fundi which appear to increase with age. Hypopigmentation was evident in the *Mitf*^*mi-vga9/+*^ heterozygotes in all age groups studied. These results suggest a progressive, age-related reduction of pigmentation and increase in depigmented spots in the fundi of these animals, most likely due to accumulation of lipofuscin debris.

**Figure 2.**
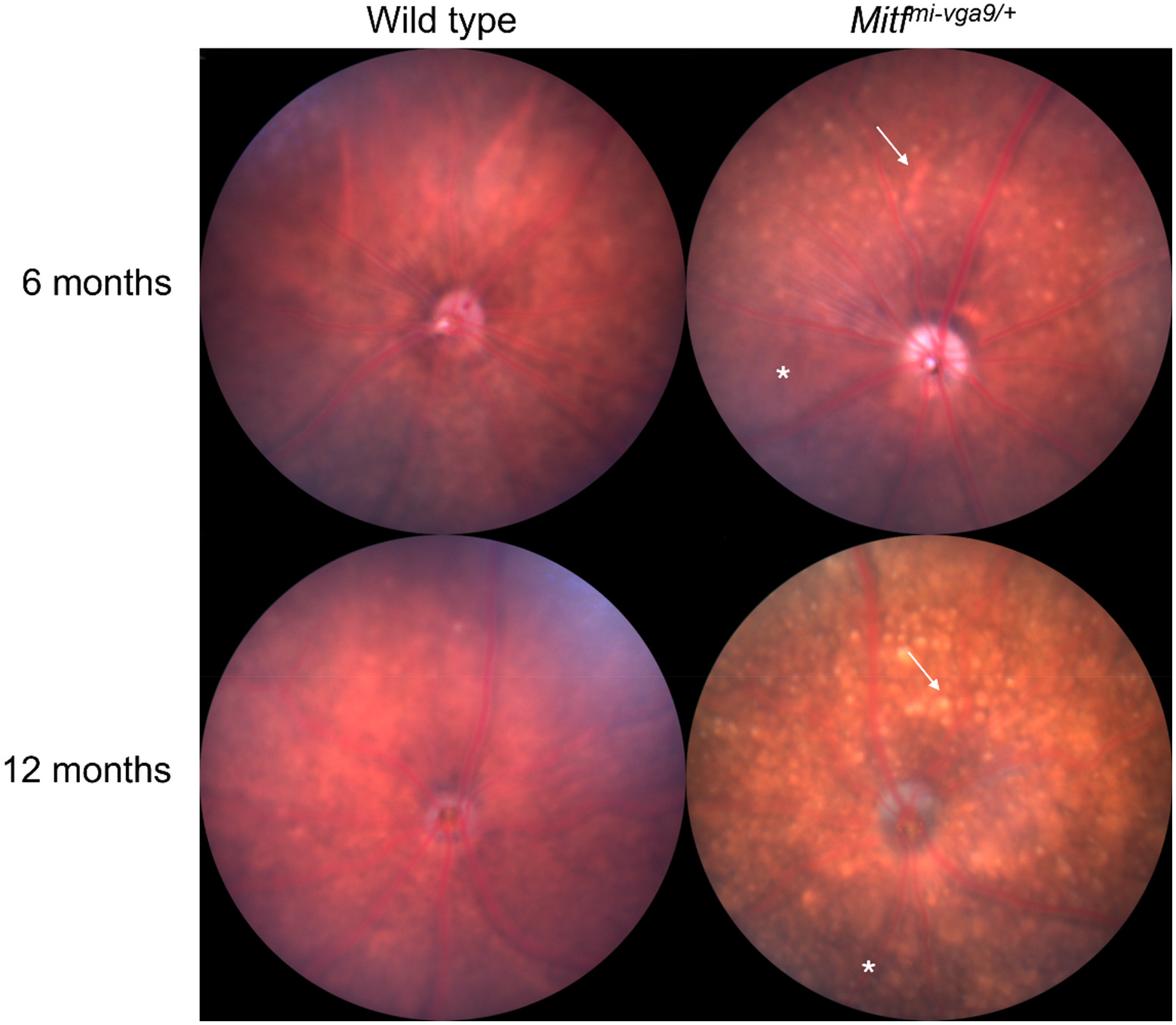
Accumulation of yellow spots increases with age in *Mitf*^*mi-vga9/+*^ mice. Representative fundus photographs from wild type and *Mitf*^*mi-vga9/+*^ mouse eyes. N=6 animals per genotype. Arrows: yellow lesions; asterisks: hypopigmentation.

To determine whether the lipofuscin build up occurred in the RPE of the *Mitf*^*mi-vga9/+*^ mice we investigated RPE flat mount preparation, using the 488 nm confocal microscope laser as the excitation source. The pigment autofluoresces at this wavelength thus allowing the measurement of the lipofuscin-like granules. 3-month-old *Mitf*^*mi-vga9/+*^ mice exhibited no signs of autofluorescence from lipofuscin accumulation, despite the presence of white spots in their fundus^29^. It is therefore possible that the depigmented spots observed in the fundi do not represent fully developed lipofuscin-like granules. Nevertheless, at 6 months of age the RPE of the *Mitf*^*mi-vga9/+*^ mice presented a clear indication of fluorescent pigment granules, which progressively build up during the aging process. At 12 months of age these fluorescent pigment granules were markedly more abundant in *Mitf*^*mi-vga9/+*^ mice than in wild type animals of the same age (Figure 3). Taken together these results indicate that lipofuscin-like granules accumulate quickly with age in *Mitf*^*mi-vga9/+*^ mice.

**Figure 3.**
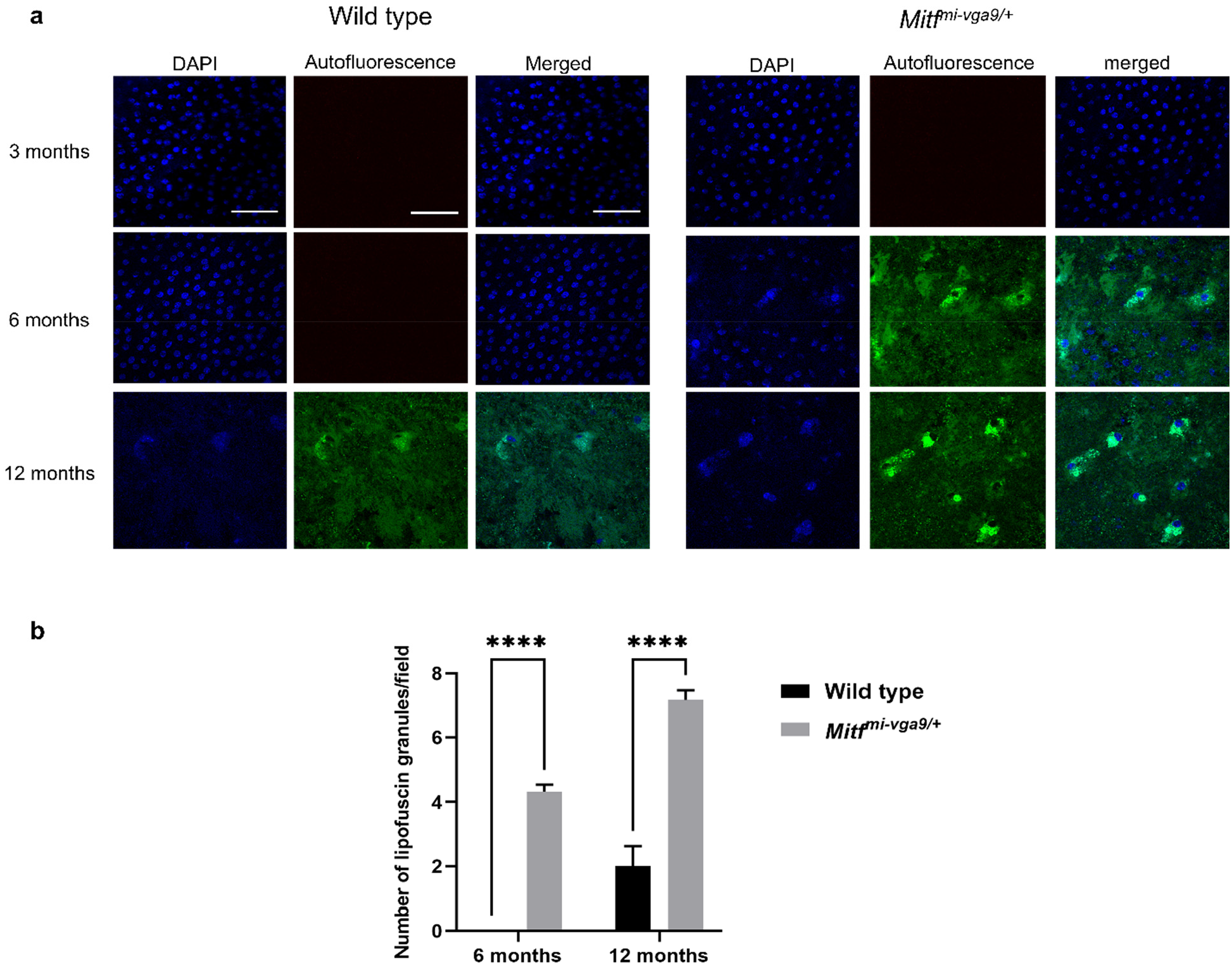
Lipofuscin accumulates faster in *Mitf*^*mi-vga9/+*^ mice than in wild type mice. (**a** and **b**) Representative images of RPE flat mounts from wild type and *Mitf*^*mi-vga9/+*^ mutant mice at 3, 6 and 12 months of age; Autofluorescence images; DAPI nuclear staining is shown in blue. Lipofuscin accumulation is indicated in green and corresponds to intrinsic autofluorescence detected in the 488 nm excitation channel under identical acquisition settings. (**c**) Quantification of fluorescent lipofuscin granules from wild type and *Mitf*^*mi-vga9/+*^ mice at 6 and 12 months of age. The number of fluorescent lipofuscin-like granules was counted in three random regions per eye. The images used for counting were all the same size and same magnification. ****P<0.0001. The values on the graph are mean ± SEM. N= 6 animals per genotype. P value was calculated using Student’s t-test. Scale bar indicates 50µm and applies to all panels.

### The RPE structure and integrity in *Mitf*^*mi-vga9/+*^ mice at 3 months of age

The RPE flat mounts from *Mitf*^*mi-vga9/+*^ animals, were labeled with phalloidin and an antibody against zonula occludens-1 (ZO-1), to determine the RPE’s structural integrity and morphology. Phalloidin binds to actin filaments and can also be utilized to detect physical injury to the RPE’s cell organization. As shown in Figure 4a, the RPE monolayers found in wild type and *Mitf*^*mi-vga9/+*^ mice presented a rather homogeneous formation of polygonal cells, containing mostly mononucleated and binucleated cells. In RPE monolayers of both wild type and *Mitf*^*mi-vga9/+*^ mice, we observed uniform ring-like microfilament networks, outlining the apical edges of separate epithelial cells. Examination of phalloidin-stained RPE flat mounts from *Mitf*^*mi-vga9/+*^ eyes indicated an absence of physical damage, with cell membranes clearly delineated and the epithelium’s hexagonal structure preserved (Figure 4a). The protein ZO-1, which is involved in membrane adherence junctions, helps in the maintenance of the integrity of the blood-retinal-barrier junctions and acts as an indicator of RPE functionality and structure. ZO-1 antibody staining of RPE flat mounts showed that wild type and *Mitf*^*mi-vga9/+*^ RPE cells were composed of a monolayer of closely arranged cells, characterized by a highly polarized epithelial morphology (Figure 4b). Both mononucleated and binucleated cells were observed in wild type and *Mitf*^*mi-vga9/+*^ RPE flat mounts, with no obvious alterations in overall epithelial organization. No apparent differences in RPE cell morphology or size were evident between genotypes. Collectively, these findings indicate that *Mitf*^*mi-vga9/+*^ mice display a regular, well-organized RPE architecture and typical barrier function.

**Figure 4.**
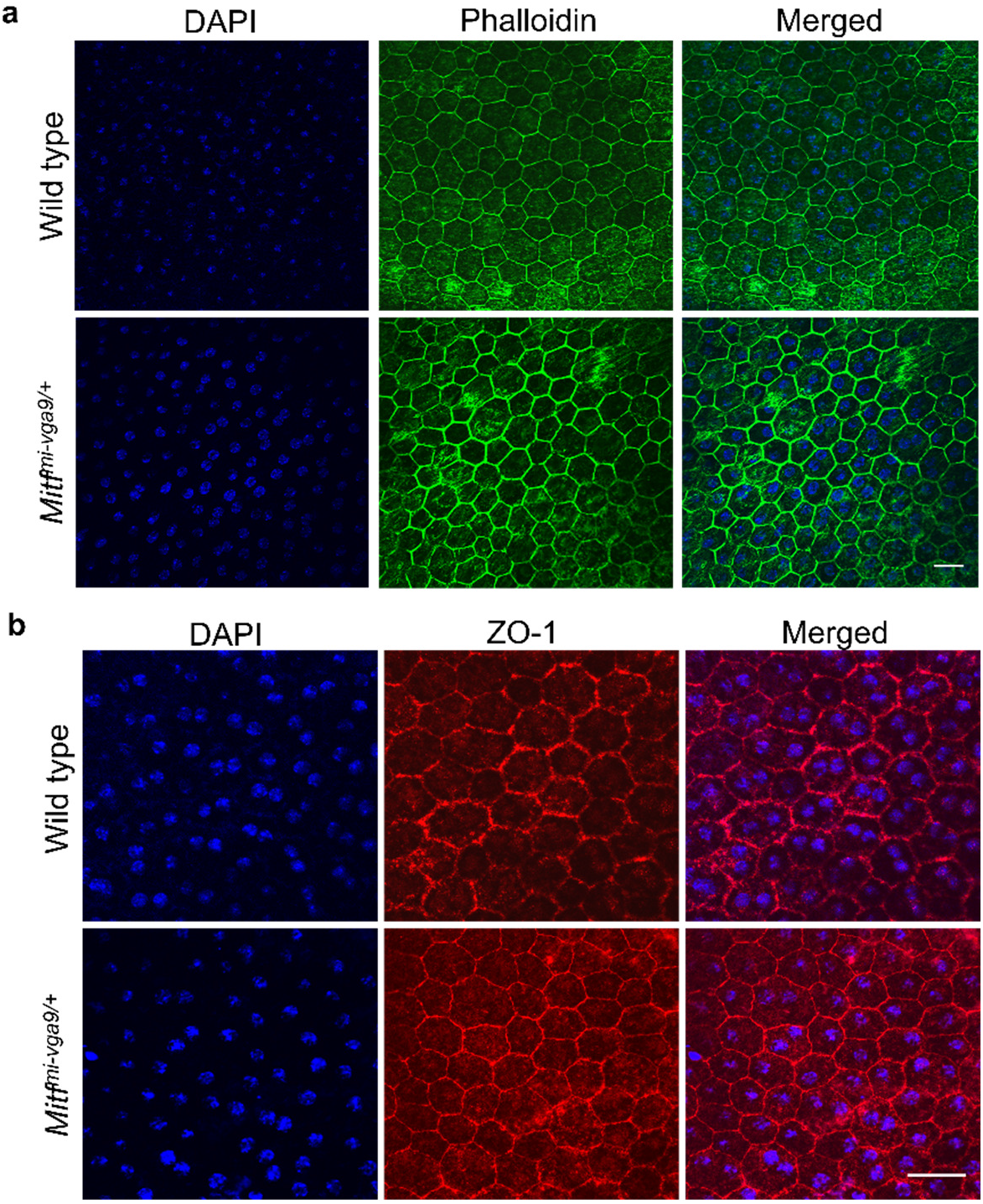
The structure and integrity of the RPE in 3-month-old *Mitf*^*mi-vga9/+*^ mice. (**a** and **b**) Representative z-stack images of RPE flat mounts. (**a**) Phalloidin immunostaining (green). (**b**) ZO-1 immunostaining (tight junction) (red). DAPI nuclear staining is shown in blue. N=6 animals per genotype. Scale bar indicates 50µm.

### *Mitf* regulates autophagy in RPE cells

Next, we examined whether the *Mitf*^*mi-vga9/+*^ mutation affects autophagy flux in primary RPE cells. Primary RPE cultures were independently generated from wild type and *Mitf*^*mi-vga9/+*^ mice and subjected to basal or starvation conditions in the presence or absence of bafilomycin A1 (Baf). Protein levels were analyzed by western blotting in four independent experiments. Autophagy flux reflects the balance between autophagosome formation and lysosomal degradation and was assessed by analyzing p62 and LC3B-II protein levels under four conditions: basal, Baf and starvation with or without Baf. Under basal conditions, *Mitf*^*mi-vga9/+*^ RPE cells displayed significantly higher levels of p62 together with increased LC3B-II levels compared to wild type cells, indicating accumulation of autophagy-related markers in the absence of treatment (Figure 5a-c). These findings are consistent with the increased *Lc3b* expression detected in eyecup tissues from 3-month-old *Mitf*^*mi-vga9/+*^ mice. In wild type RPE cells, Baf treatment led to the expected accumulation of p62 and LC3B-II, consistent with active autophagic flux. In mutant RPE cells, p62 levels further increased following Baf treatment, whereas LC3B-II levels did not show a significant additional accumulation, suggesting impaired autophagic clearance. Starvation induced an increase in LC3B-II protein levels in both genotypes, reflecting enhanced autophagosome formation. Under starvation conditions, Baf treatment resulted in a clear additional accumulation of LC3B-II and p62 in wild type RPE cells. In contrast, the LC3B-II response to Baf treatment was markedly blunted in *Mitf*^*mi-vga9/+*^ RPE cells, while p62 levels remained significantly elevated regardless of Baf treatment (Figure 5b and c). Taken together, the elevated basal levels of LC3B-II and p62, combined with the blunted LC3B-II response upon lysosomal inhibition under both basal and starvation conditions, indicate that autophagosomes accumulate in *Mitf*^*mi-vga9/+*^ RPE cells due to defective autophagic degradation rather than increased autophagic induction. These results support the presence of impaired autophagy flux in mutant RPE cells.

**Figure 5.**
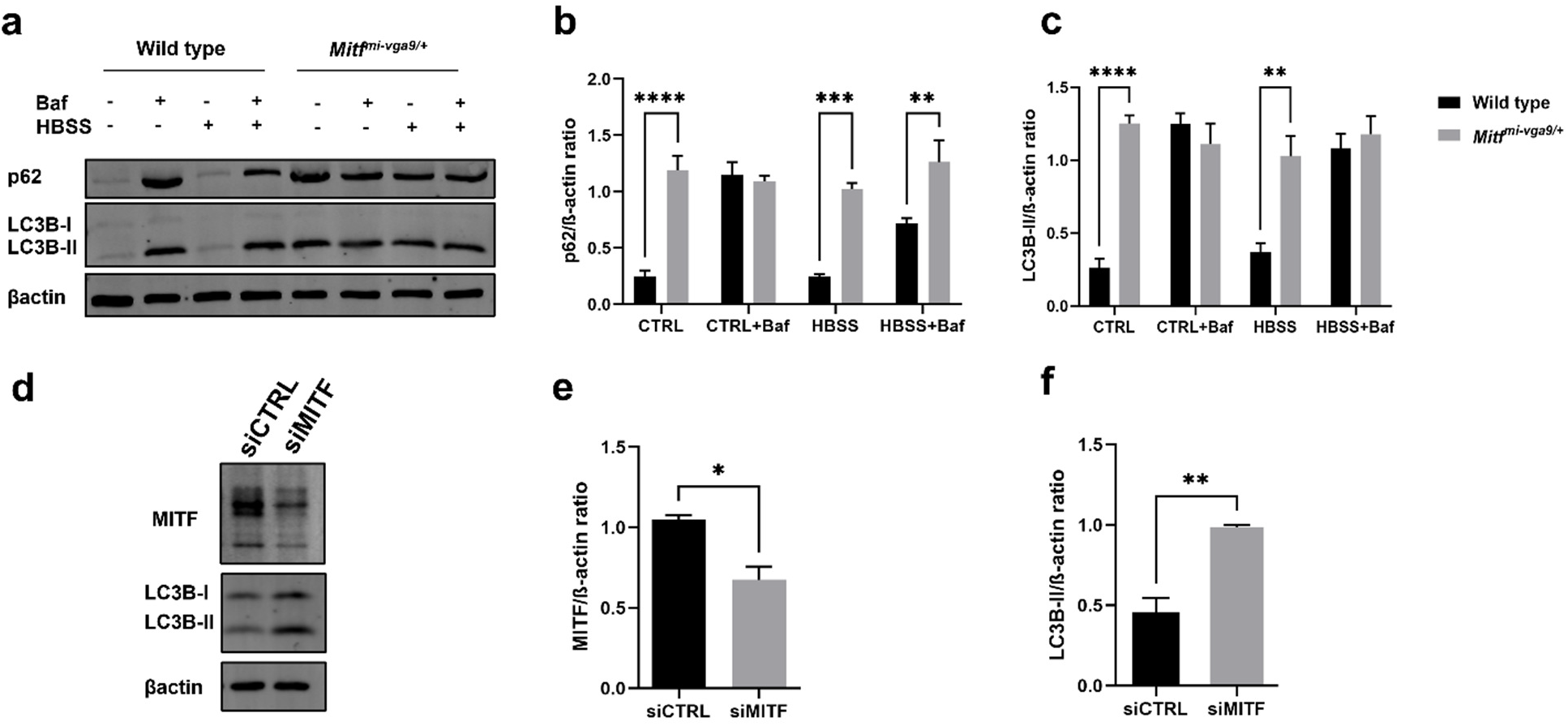
Impaired autophagy flux in *Mitf*^*mi-vga9/+*^ mice. (**a-c**) Western blot analysis of normal and mutant mouse primary RPE cells in normal growth media or starvation media (HBSS) with or without Baf-A1. The graphs show quantification of four independent experiments. The western blots show a representative image. Proteins were probed using antibodies against, p62, LC3B-II and βactin as a loading control. (**a**) Representative Western blot image from four independent experiments. (**b** and **c**) Mean quantification of the protein expression of p62 and LC3B-II normalized to β-actin from four independent experiments. **P< 0.01, ***P< 0.001 and ****P< 0.0001. The values on the graphs are mean ± SEM. P-values were calculated using the two-way ANOVA followed by Šidák post hoc multiple comparisons test. (**d-f**) Western blot analysis of protein lysates from siMITF and siCTRL treated ARPE-19 cells in normal medium. (**d**) The blots were stained with anti-MITF and anti-LC3B antibodies. The figure is representative of three independent experiments. (**e** and **f**) Quantification of the intensity of MITF and LC3B-II bands normalized to β-actin. *P< 0.05, **P< 0.01. The values on the graphs are mean ± SEM. P-values were calculated using Student’s t-test.

To determine if MITF regulates autophagy in human RPE, we performed experiments using the human ARPE-19 cell line. Knockdown of MITF resulted in approximately 50% reduction in MITF protein expression compared to siCTRL (5d-f). In addition, we observed a significant increase in basal levels of LC3B-II in ARPE-19 cells treated with siMITF compared to siCTRL. This is therefore consistent with our results obtained using primary RPE cells. Elevated LC3B-II expression might indicate either increased autophagic activity or diminished clearance of autophagosomes. These observations suggest that MITF is crucial in modulating autophagy in mouse primary RPE cells and human ARPE-19 cells.

Mature autolysosomes (ALs) were further analyzed by electron microscopy obtained from the RPE of 3-month-old wild type and *Mitf*^*mi-vga9/+*^ mice. ALs were identified based on established ultrastructural features, characterized by single-membrane vesicles containing electron-dense and partially degraded material. The ALs were quantified under blinded conditions; the investigator remained unaware of the experimental group assignments during both image acquisition and analysis. In each sample, ALs were quantified across four randomly chosen areas within every cell. The images from which these counts were derived were consistent in terms of dimensions and magnification. As illustrated in Figure 6a, *Mitf*^*mi-vga9/+*^ animals displayed: lack of pigmentation, hypotrophy and increased presence of ALs. We found a marked difference in the quantity of ALs within the mutant compared to the control RPE (Figure 6b). These results indicate that impaired autophagy in *Mitf*^*mi-vga9/+*^ RPE causes a buildup of non-functional ALs.

**Figure 6.**
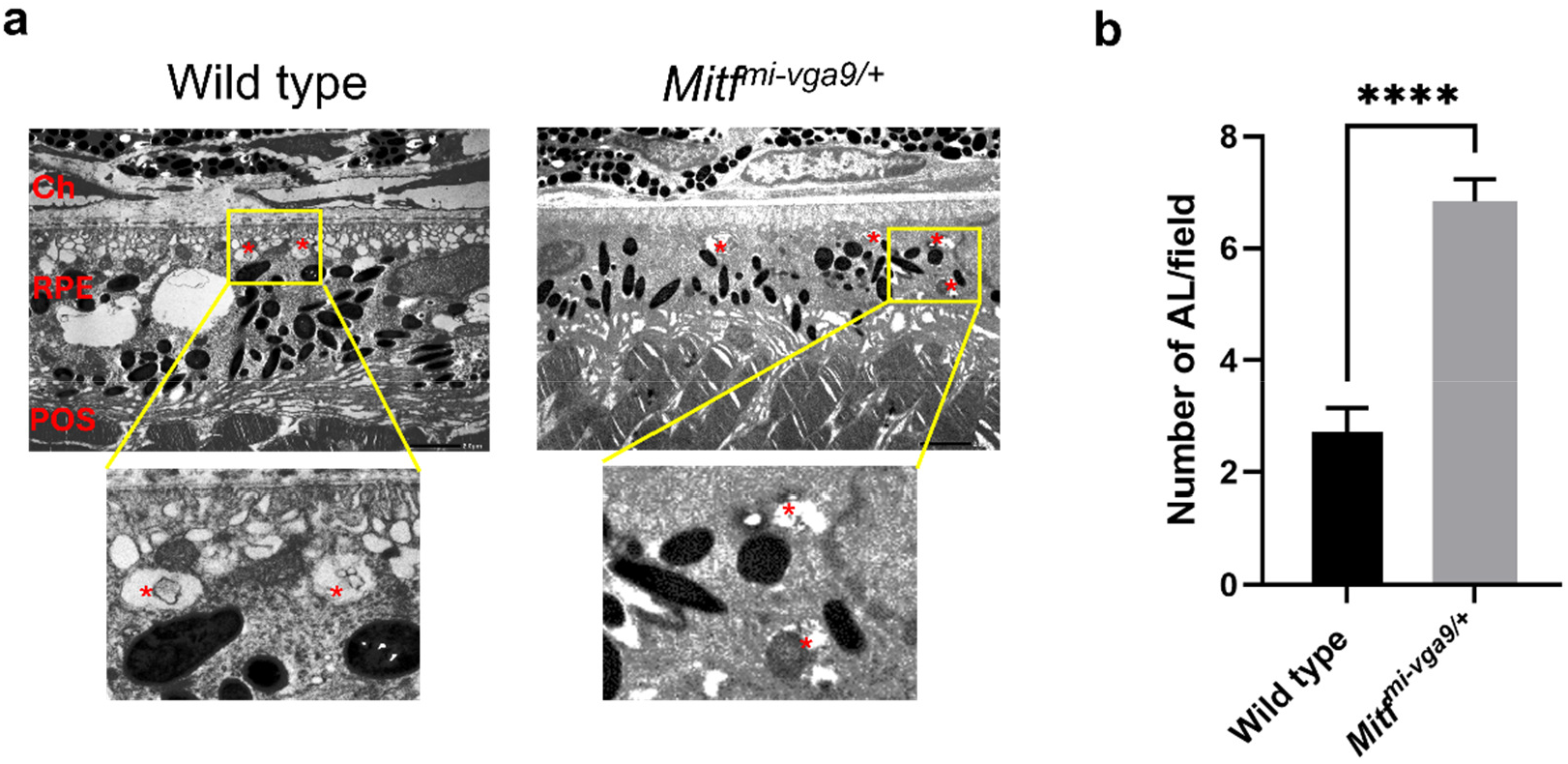
The number of autolysosomes in RPE cells from 3-months-old wild type and *Mitf*^*mi-vga9/+*^ mice. (**a**) EM images from normal and mutant mice respectively. (**b**) Autolysosomes, as identified as single-membrane vesicular structures containing electron-dense and partially degraded cytoplasmic material, in contrast to double membrane autophagosomes, were counted in three random regions per cell. Ch, choroid; RPE, retinal pigment epithelium; POS, photoreceptors. Red asterisks: autolysosomes. All images used for quantification were acquired at the same magnification and field size. N=6 animals per genotype. Scale bar indicates 2 µm. ****P< 0.0001. The values on the graph are mean ± SEM. P values were calculated using Student’s t-test.

## Discussion

In this study, we demonstrate that *Mitf* is a crucial regulator of autophagy in the RPE in vivo. Using the *Mitf*^*mi-vga9/+*^ mouse model, we show that reduced *Mitf* function leads to impaired autophagic flux in RPE cells, characterized by accumulation of LC3B-II and p62, increased numbers of autophagosomes and autolysosomes, and transcriptional dysregulation of key autophagy- and lysosome-related genes. Importantly, these autophagic defects precede overt structural alterations of the RPE and are associated with a progressive age-dependent accumulation of lipofuscin-like granules. Consistent with that results obtained in human RPE cells further support a conserved role for MITF in autophagy regulation. However, the structure and integrity of the mouse RPE was found to be unaffected by the *Mitf*^*mi-vga9/+*^ mutation. The timing of accumulation of lipofuscin-like granules is affected by the *Mitf*^*mi-vga9/+*^ mutation, in that they appear earlier and faster in the RPE of the mutant mice than in wild type mice. Taken together, our findings support a role for *Mitf* in regulating RPE autophagy and lysosomal homeostasis, providing insight into mechanisms that govern long-term RPE maintenance and intracellular waste handling. Recent work has shown that loss of MITF can alter TOR-pathway gene expression and promote nuclear TFE3 localization, suggesting that some of the autophagy-related alterations observed here may also involve secondary changes in MiT/TFE signaling pathways rather than solely direct effects of MITF deficiency^30^.

Analysis of gene expression in the RPE-choroid complex in *Mitf*^*mi-vga9/+*^ mice revealed age-dependent alterations in the transcription of autophagy- and lysosome-associated genes (Figure 1). Reduced expression of the *Mitf* itself was consistently observed in mutant animals across all ages examined ^19,28^, with expression levels reduced substantially beyond those expected from a simple heterozygous effect. The transcriptional changes observed in *Mitf*^*mi-vga9/+*^ mice were consistent with a broader role of MITF in the regulation of lysosomal and autophagy-related gene networks. MITF belongs to the MiT-TFE family of transcription factors, which has been extensively implicated in the control of lysosomal biogenesis and autophagy through coordinated regulation of degradative pathways^12,31,32^. In this context, the increased expression of *Lc3b* together with the reduced expression of *Lamp1* suggests a dysregulation of the autophagy-lysosome system rather than isolated changes in individual components^33,34^. However, changes in the expression of autophagy-related genes do not necessarily predict functional autophagy activity, underscoring the importance of assessing autophagy flux directly^35,36^. Previous studies have shown that MITF regulates autophagy-lysosome pathways in other cellular contexts, including melanoma cells^14,37^, supporting a conserved role for MITF in degradative pathway regulation. Overall, our transcriptional data provide a mechanistic framework that is functionally substantiated by the impaired autophagy flux and ultrastructural alterations observed in RPE cells.

Taken together, the fundus (Figure 2) and confocal imaging data (Figure 3) indicate that the early fundus appearance changes observed in *Mitf*^*mi-vga9/+*^ animals are unlikely to reflect lysosomal waste accumulation. Although discrete white spots are already present at 3 months of age, lipofuscin-like granules are not detected until later stages, suggesting that the initial funduscopic alterations arise from mechanisms distinct from lysosomal storage^29^. By contrast, 6-month-old *Mitf*^*mi-vga9/+*^ mice display increased accumulation of lipofuscin-like granules compared to age-matched wild type mice, and this accumulation further increases by 12 months of age. The delayed emergence of these autofluorescent granules is consistent with impaired autophagy-lysosome dynamics in mutant RPE cells. Reduced *Lamp1* expression, together with the accumulation of ALs, suggests that lysosomal maintenance and degradative efficiency are compromised, limiting the capacity of RPE cells to process intracellular cargo over time^33,34^. Under such conditions, incompletely degraded material may persist within the lysosomal compartment and progressively accumulate as lipofuscin. This interpretation is supported by previous studies indicating that impaired lysosomal function and defective autophagic flux favor the buildup of non-degradable autofluorescent material in long-lived cells^38,39^. Reduced *Mitf* expression may influence genes involved in RPE homeostasis and retinoid metabolism, including *Rpe65* and *Abca4*, both of which have been associated with lipofuscin accumulation and retinal degeneration^39–43^. Together, these findings support a potential connection between altered retinal homeostasis and the accumulation of lipofuscin-like material observed in *Mitf*^*mi-vga9/+*^ mice.

Impaired autophagy flux observed in *Mitf*^*mi-vga9/+*^ primary RPE cells provides functional evidence that reduced *Mitf* activity disrupts late stages of the autophagic pathway (Figure 5). Accumulation of LC3B-II and p62 proteins under basal and starvation conditions, together with the absence of further increases upon lysosomal inhibition, is widely interpreted as indicative of defective autophagic degradation rather than enhanced autophagosome biogenesis^44–46^. Consistent with the transcriptional role of MITF discussed above, previous studies in melanoma cells have shown that MITF influences autophagy-lysosome pathways, providing a mechanistic framework that supports the functional defects in autophagy flux observed here^14,31^. Consistent with a late-stage defect, ultrastructural analyses in our model revealed an increased number of ALs, suggesting that autophagosome-lysosome fusion occurs but that terminal cargo degradation is inefficient^47^ (Figure 6). Importantly, reduction of MITF in human ARPE-19 cells resulted in comparable increases in LC3B-II protein levels, indicating that this regulatory role of MITF on autophagy is conserved beyond melanocytic lineages (Figure 5f). Together, these findings indicate that *Mitf* insufficiency compromises autophagy flux at a late degradative step in RPE cells.

In conclusion, this study supports an important role for *Mitf* in the long-term maintenance of autophagy-lysosome function in the RPE. By integrating in vivo, ex vivo, and human cell-based analyses, we have shown that reduced *Mitf* activity leads to impaired autophagy flux, accumulation of ALs, and a delayed build up of lipofuscin-like granules, collectively reflecting a progressive decline in intracellular degradative capacity. These findings highlight *Mitf* as a key regulator of RPE homeostasis and provide a mechanistic framework linking transcriptional control of lysosomal pathways to age-dependent cellular dysfunction.

## Material and methods

### Animals

Wild type (C5BL/6J) and *Mitf*^*mi-vga9/+*^ mutant mice were used in this study. All the mouse strains bred and raised at the animal facilities at the Biomedical Center, University of Iceland. The mice were kept transparent polypropylene cages with wire tops (Techniplast, Italy). They were maintained on a 12 h light and dark cycle, at 21-22°C ambient temperature and 60% humidity. The animals had free access to food and water. All the experiments on mice were approved by the Icelandic and Food Veterinary Authority (MAST), license no. 2017-04-03. Additionally, all the animals were treated in line with the Association for Research in Vision and Ophthalmology (ARVO) Statement for Use of Animals in Ophthalmic and Vision Research.

### Fundus photography

In vivo fundus images were acquired from anesthetized mice using a Micron IV rodent fundus imaging system (Phoenix Research Labs Inc., Pleasanton, CA, USA). Before the funduscopic examination, the mice received an intraperitoneal administration of a solution composed of 40 mg/kg^−1^ Ketamine (Pfizer, Denmark) and 4 mg/kg^−1^ Xylazine (Chanelle Pharmaceutical, Ireland) and the anesthesia was continued, administered at a fifty-percent reduction, according to requirements. Topical application of Mydriacyl (1% tropicamide) (Alcon Inc., USA) eyedrops facilitated pupil dilation, and Alcaine (Alcon Inc., USA) drops were subsequently utilized to provide corneal anesthesia. Concurrent application of a 1% methylcellulose gel and a 2.5mm diameter mini contact lens (Ocuscience LLC, Henderson, NV) to the cornea of both eyes was performed to minimize the incidence of cataract formation^48,49^. In preparation of fundus photography, the contact lens of one eye was displaced, and thereafter, 1% methylcellulose gel was reapplied to the corneal surface of the eye. Bright field images with the optic disc in the center of the image were captured as previously described^29^.

### RPE flat mounts

Wild type and *Mitf*^*mi-vga9/+*^ mice at 3, 6 and 12 months of age were sacrificed by cervical dislocation. The eyes were enucleated and fixed in fresh 4% paraformaldehyde (PFA) in PBS for 2 h. The anterior segments of the eyes (cornea, iris, ciliary body and lens) were removed. Retinal tissue was carefully removed from the eyecup, and the remaining cups containing RPE, choroid and sclera were dissected into quarters by radial cuts and washed 3 times in PBS for 5 min. The RPE/choroid flat mounts were permeabilized in PBS with 0.1% Triton X-100 for 20 min and then blocked with 10% normal goat serum, 0.5% Triton X-100 in PBS for 1h. Next, the samples were stained with Alexa Fluor 488 phalloidin (1:500, Invitrogen) to visualize the cytoskeleton and with ZO-1 antibody (1:200, ZO1-1A12, Invitrogen) to stain cell boundaries. The antibodies were diluted in blocking buffer solution overnight at 4°C, or the tissue was incubated with blocking buffer to visualize accumulation of lipofuscin-like granules accumulation by detecting intrinsic autofluorescence using 488 nm excitation. After three washes in PBS (5 min per wash), the RPE-choroid complexes were incubated with the appropriate secondary antibody in blocking buffer solution for 1h at room temperature (RT). The secondary antibody used was goat anti-mouse IgG H&L Alexa Fluor 546 (1:1000, Life Technologies). After 3 washes in PBS (5 min each), the RPE/choroids were flat-mounted on glass slides and cover slipped with the Fluoroshield mounting medium containing 4′,6-diamidino-2-phenyllindole (DAPI; ab104139, Abcam). The images were acquired with the Olympus FV10-MCPSU confocal microscope. The number of nuclei of lipofuscin-like granules bodies were quantified in RPE flat mounts from six animals per genotype by the “blinded” observer in four randomly selected areas and averaged using ImageJ software.

### RPE primary cell culture

Three-month-old wild type and *Mitf*^*mi-vga9/+*^ mice were sacrificed by cervical dislocation. Intact eyes with optic nerves were gently removed using curved scissors and immediately dipped in a 3.5 cm Petri dish containing PBS and 1% penicillin/streptomycin (Thermo Fischer Scientific) on ice. Eyes were then washed with PBS. With the help of a dissecting stereomicroscope, all the muscles and tissues around the eyeballs were carefully removed using forceps and angled scissors, and then washed again to clean away the remaining blood and extra tissue before placing the eyeballs in 2% Dispase (Thermo Fischer Scientific) in DMEM-F12 medium for 45 min at 37°C and in 5% CO_2_. The eyeballs were then transferred to a dish with growth medium consisting of DMEM (1x) high glucose, 10% fetal bovine serum (FBS) and 2% penicillin/streptomycin (all reagents purchased from Thermo Fischer Scientific). Next, the eyeballs were placed in a new petri dish with PBS for dissection. Under a dissecting microscope, the eyes were cut along the corneal sclera edge. The cornea, iris epithelium and lens were gently removed. After removal of the retina, the RPE and the posterior sclera were put in 0.25% trypsin (Thermo Fischer Scientific) and incubated for 30 min at 37°C (and 5% CO_2_) to help the separation of the RPE from the sclera. After the incubation, a single layer of RPE cells can be clearly observed under the microscope at this time. RPE cells were placed in a 15 mL falcon tube prior to centrifugation at 1300 rpm for 5 min at RT. The supernatant was discarded and then the RPE cells were gently resuspended in 2 mL of growth medium. As some of the RPE fragments were large, a p200 micropipette was used to pipette the sheets gently about 10 times. RPE cells isolated from both eyes of a single wild type or *Mitf*^*mi-vga9/+*^ mouse were pooled and plated into one well of a 6-well plate containing 500 µL of growth medium. Cells were confirmed to be evenly distributed under a microscope before incubating at 37°C in 5% CO_2_ incubator. The culture dish was kept in undisturbed condition for at least 48h to allow cell attachment to the dish. Every 2 days, half of the culture medium was removed and replaced with fresh medium. Primary cultures within the first three to five passages were used for the experiments.

### Primary RPE cell treatment

Primary cultured cells were maintained in DMEM high glucose supplemented with 10% of FBS and 1% of penicillin-streptomycin (10000 U/mL) or incubated in HBSS (Thermo Fisher Scientific) as a starvation medium, in the presence or absence of Bafilomycin A1 (Baf; Santa Cruz) at 100 nM. After 24 h in culture, the cells were subjected to the indicated conditions for 24 h prior to protein extraction.

### RNAi treatment

ARPE-19 cell were cultured in a 6-well plate 24 h and then transfected with 50nM of the appropriate siRNA for 72 h before protein extraction. Lipofectamine RNAiMax (Thermo Fisher Scientific, #13778-075) and siRNA reagents were added to Opti-MEM medium (Thermo Fisher Scientific, #31985062) according to manufacturer’s protocol. The siRNAs used were siCTRL (Thermo Fisher Scientific, #1299099) and siMITF (Thermo Fisher Scientific, #HSS142939).

### Protein extraction and western blotting

Mouse primary RPE cells from wild type and *Mitf*^*mi-vga9/+*^ mice were cultured in 6 well plates and lysed with RIPA buffer with 1:100 PMSF (100mM; Sigma-Aldrich), DTT (Thermo Fisher Scientific), 1:100 Protease Inhibitor Cocktail (Sigma-Aldrich) and 1:6 Laemmli buffer. The lysate was centrifuged at 14000g for 20 min at 4ºC. Finally, the supernatant was boiled at 95 ºC for 5 min. The samples were then run on a 12.5% SDS-PAGE gels and transferred onto a 0.2 µm PVDF membrane (Thermo Fisher Scientific). The membranes were blocked in 5% bovine serum albumin (BSA; Sigma-Aldrich) in TBS-T for 1h at RT followed by overnight incubation at 4°C with the appropriate primary antibodies diluted in blocking buffer: rabbit anti-LC3B (1:100, 2775S, Cell Signaling), rabbit anti-p62 (1:1000, 5114S, Cell Signaling), mouse anti-MITF C5 (1:1000, ab12039, Abcam) and rabbit anti-β-actin (4970S, 1:2000, Cell Signaling) as a load control. Membranes were washed three times with TBS-T and incubated with the appropriate secondary antibody for 45 min at RT diluted in blocking buffer. The secondary antibodies used were anti-mouse IgG(H+L) Dylight 800 conjugate (1:10000) and anti-rabbit IgG(H+L) Dylight 680 conjugate (1:10000) from Cell Signaling. The images were captured using Odyssey CLx Imager (LI-COR Biosciences). To quantify protein bands, the digital images obtained from the Odyssey scans as bands on the western blots, were quantified using ImageJ and normalized to actin levels.

### Electron microscopy

Three-month-old wild type and *Mitf*^*mi-vga9/+*^ mice were sacrificed by cervical dislocation. Enucleated whole eyes were immediately fixed in a 2.5% glutaraldehyde solution (Electron Microscopy Sciences) in phosphate buffer (PB) (0.1 M, pH 7.4) (Sigma-Aldrich) over night at 4ºC. Ocular tissues were then post fixed with a fresh mixture of 2% osmium tetroxide (Electron Microscope Sciences) with 1.5% potassium ferrocyanide (Sigma-Aldrich) in PB for 1 h at RT. The samples were then rinsed two times in PB and dehydrated in graded concentrations of an ethanol solution (Sigma Aldrich) (50% for 5 min, 70% ethanol for 5 min, 4% uranyl acetate in 70% for 15 min, 80% for 5 min, 90% for 5 min and 96% once for 5 min, once for 10 min and twice for 15 min). This was followed by infiltration of ethanol and Supurr’s Resin for 1 h at 1:1 dilution. The mixture was removed and replaced with pure resin and rotated for 1 h. The capsules were filled with resin which was allowed to polymerize over night at 70ºC in an oven. Ultrathin sections (70 nm) were cut transversally on a Leica Ultracut microtome (Leica Microsystems), stained with toluidine blue and picked up on a 2-by 1-mm nickel slot grid (Electron Microscopy Sciences) coated with polystyrene film (Sigma-Aldrich). Sections were post stained with 4% uranyl acetate (Sigma-Aldrich) in H_2_O for 10 min, rinsed several times with H_2_O (Sigma-Aldrich) followed by incubation in 0.4% Reynolds’ lead citrate (ref) in H_2_O for 10 min and rinsed several times with H_2_O. Micrographs were taken with a JEM-1400Plus Transmission Electron Microscope, and the number of autolysosome counted by the “blinded” observer in 3 randomly selected areas and averaged using ImageJ software.

### Gene expression analysis by quantitative reverse transcription PCR (RT-qPCR)

Wild type and *Mitf*^*mi-vga9/+*^ mice of 3 and 12 months of age were sacrificed by cervical dislocation and the eyeballs dissected and flash frozen in liquid nitrogen. Samples were kept at −80C until RNA was isolated using the RNeasy mini kit (cat no. 74104 Qiagen) according to the manufacturer’s instructions. cDNA was produced from 500 ng of total RNA using an iScript first-strand kit (cat no. 1708891 Bio-Rad Laboratories). iQSYBR green supermix (cat no. 1708891) and certified PrimePCR oligos (MITF, cat no qMmuCID0013258; LC3B, cat no qMmuCED0004713; RAB7, cat no qMmuCID0005411; LAMP1, cat no qMmuCID0027030) were used to determine transcript expression in a CFX96 thermal cycler (all Bio-Rad Laboratories). Threshold values were analyzed by the ΔΔCt method, and data were determined relative to two housekeeping genes, TATA-box binding protein (TBP) (cat no qMmuCID0040542) and hypoxanthine phosphoribosyltransferase 1 (HPRT1) (cat no qMmuCID0005679).

### Statistical analysis

Results from three or more independent experiments are presented as means ± standard error of the mean (SEM) unless stated otherwise. Statistical analysis and graphs were plotted using GraphPad Prism v.9.0 (GraphPad Software, San Diego, CA, USA). Data were analyzed using Student’s t-test and two-way ANOVA followed by Šidák post hoc multiple comparisons test as appropriate. Differences of *P<0.05, **P<0.01, ***P<0.001 and ****P<0.001 were considered statistically significant.

## Acknowledgements

We thank Jóhann Arnfinnsson and Paulina Cherek for processing the samples and excellent technical assistance with the EM. We also thank Prof. Kai Kaarniranta for expert input on autolysosomal identification.

## Contributions

A.G-LL was involved with designing the experiments, performing the fundus photographs, RPE flat mounts, RPE primary cell culture experiments and collecting samples, analyzing data, and writing the manuscript, K.H was involved in the RNAi experiments, F.L was involved in the qPCR experiments, H.A was involved with designing the experiments, E.S was involved with designing the experiments and writing the manuscript, and provided the experimental animals, M.H.O was involved with designing the experiments, supervising the experimental work and writing the manuscript, T.E was involved with designing the experiments, supervising the experimental work and writing the manuscript.

## Data availability

All data supporting the findings of this study are available within the paper. The datasets generated and analyzed during the current study are available from the corresponding author upon reasonable request.

